# Cercosporamide inhibits bone morphogenetic protein receptor type I kinase activity in zebrafish

**DOI:** 10.1101/2020.05.29.123588

**Authors:** Jelmer Hoeksma, Gerard C.M. van der Zon, Peter ten Dijke, Jeroen den Hertog

## Abstract

Zebrafish models are well established tools for investigating underlying mechanisms of diseases. Here, we identified cercosporamide, a metabolite from the fungus *Ascochyta aquiliqiae*, as a potent bone morphogenetic protein (BMP) type I receptor kinase inhibitor through a zebrafish embryo phenotypic screen. The developmental defects in zebrafish, including lack of the ventral fin induced by cercosporamide was strikingly similar as the phenotypes caused by renowned small molecule BMP type I receptor kinase inhibitors and inactivating mutations in zebrafish BMP receptors. In mammalian cell-based assays, cercosporamide blocked BMP/SMAD-dependent transcriptional reporter activity and BMP-induced SMAD1/5-phosphorylation. Biochemical assays with a panel of purified recombinant kinases demonstrated that cercosporamide directly inhibited kinase activity of BMP type I receptors (also called activin receptor-like kinases (ALKs)). In mammalian cells, cercosporamide selectively inhibited constitutively active BMP type I receptor-induced SMAD1/5 phosphorylation. Importantly, cercosporamide rescued the developmental defects caused by constitutively active Alk2 in zebrafish embryos. Taken together, we believe cercosporamide may be the first of a new class of molecules with potential to be developed further for clinical use against diseases that are causally linked to overactivation of BMP receptor signaling, including Fibrodysplasia ossificans progressiva and diffuse intrinsic pontine glioma.

## Introduction

Zebrafish (*Danio rerio*) is an attractive model for studying biological effects of both genetic mutations and chemical compounds *in vivo*. Zebrafish are vertebrates with a highly conserved physiology that develop all organs and primary tissues in several days (Kimmel et al., 1995). Moreover, zebrafish embryos are transparent, which makes development easy to follow and defects induced by compounds or mutations easy to observe (Kimmel et al., 1995). Finally, large numbers of eggs can be obtained due to the high fecundity, making zebrafish the perfect model for genetic studies and high throughput compound screens (den Hertog, 2005; Wiley et al., 2017).

Zebrafish are frequently and intensively being used to investigate signaling in development and disease. For instance, BMP receptor signaling is widely studied in zebrafish. Bone morphogenetic proteins (BMPs) are highly conserved secreted cytokines with key roles in organ formation and tissue homeostasis (Katagiri and Watabe, 2016). Depending on dose, BMPs induce different cell fates, and control patterning within multicellular organisms during embryogenesis (Bier and De Robertis, 2015). A relatively easy to follow process in zebrafish involving BMP-signaling is dorsoventral patterning. Knock-out mutations introduced in distinct genes of the signaling cascade, including ligands, e.g. *bmp2* (Kishimoto et al., 1997), receptors, e.g. *activin receptor-like kinase* 2 (*alk2)* (Bauer et al., 2001) (in zebrafish also known as *alk8* or *lost-a-fin* (Mintzer et al., 2001)) or intracellular messengers, e.g. *smad5* (Kramer et al., 2002), all result in a dorsalization phenotype, including promotion of dorsal ectodermal cell fates at the expense of ventral tissues. Conversely, excess signaling caused by overexpression of Bmps, such as Bmp2 or Bmp7 (Schmid et al., 2000) or loss of secreted Bmp antagonists such as Chordin, Follistatin and Noggin (Dal-Pra et al., 2006), causes ventralization, the expansion of ventral tissue at the expense of dorsal structures. Overexpression of constitutively active Alk2 causes severe ventralization, whereas *alk2* null-mutants show the opposite phenotype of dorsalization and the loss of the ventral fin (Shen et al., 2009).

Overactive BMP signaling has been implicated in a large plethora of human diseases. The most prominent example is the rare genetic disorder Fibrodysplasia Ossificans Progressiva (FOP) in which fibrous tissue, such as muscles and ligaments are progressively replaced by bone tissue (Pignolo and Kaplan, 2018). FOP patients carry a missense mutation in the gene encoding the BMP receptor, activin A receptor type 1 (ACVR1, also known as ALK2) (Kaplan et al., 2009). The mutation in *ALK2* results in a gain of function of ALK2. Despite great effort in recent years, there is currently no approved treatment for FOP (Pignolo and Kaplan, 2018). Moreover, *ACVR1* is also found mutated in about 25% of patients with the rare childhood brainstem tumor diffuse intrinsic pontine glioma (DIPG) (Taylor et al., 2014). In addition, more common diseases such as myeloid leukemia (Lefort and Maguer-Satta, 2020), chronic kidney disease (Kajimoto et al., 2015), vascular calcification (Derwall et al., 2012) and atherosclerosis (Saeed et al., 2012) are also linked to overactive BMP signaling. Zebrafish have been used to study the causal involvement of BMP-signaling in a variety of skeletal and ocular diseases (Ye et al., 2009), including congenital FOP (LaBonty and Yelick, 2018; Mucha et al., 2018), radioulnar synostosis (Suzuki et al., 2020) and superior coloboma (Hocking et al., 2018). Targeting overactive BMP signaling for therapeutic gain has promise, but will require selective intervention.

BMPs exert their multifunctional effects on cells by interacting with selective cell surface BMP type I and type II receptors that are endowed with intracellular serine/threonine kinase domains. The type I receptors are also termed activin receptor-like kinases (ALKs). Upon BMP-induced type I/type II heteromeric complex formation, the constitutively active type II kinase phosphorylates the type I receptors on particular serine and threonine residues (Gomez-Puerto et al., 2019). Activated type I receptors promote phosphorylation of receptor-regulated SMAD1, SMAD5 and SMAD8 proteins, which act as transcription factor complexes by partnering with SMAD4. These heteromeric SMAD complexes translocate into the nucleus where they interact in a DNA sequence dependent manner with enhancers/promoters of target genes and regulate their expression (Hill, 2016). A typical target gene is ID1, and multimerizing the SMAD1/5 response elements in front of a minimal promoter generates a highly selective reporter system to interrogate BMP/SMAD signaling (Korchynskyi and Ten Dijke, 2002). All 4 BMP type I receptors (ALK1, ALK2, ALK3 and ALK6) activate the SMAD1/5 pathway. Ectopic expression of mutant, constitutively active BMP type I receptors mimic the BMP signaling response. Type I receptors determine the signaling specificity in BMP-induced heteromeric complexes (Gomez-Puerto et al., 2019).

BMP inhibitors have been identified by small molecule compound screens using zebrafish embryos. The first inhibitor to be identified was dorsomorphin, which induces developmental defects that phenocopy BMP-mutants (Yu et al., 2008). Dorsomorphin targets type 1 BMP-receptors, and moreover, rescues the phenotype caused by overexpression of constitutively active Alk2 in zebrafish (Shen et al., 2009; Yu et al., 2008). Unfortunately, dorsomorphin also harbors off-target effects, such as inhibition of vascular endothelial growth factors (Vegf) and AMP-activated protein kinase (Ampk) (Cannon et al., 2010; Hao et al., 2010; Zhou et al., 2001). However, related BMP-inhibitors such as DMH-1 and LDN-193189, sharing the same pyrazolo[1,5-a]pyrimidine core as dorsomorphin, have less off-target effects and are potently targeting type 1 BMP receptor kinases, predominately Alk2 (Hao et al., 2010). Finally, more recent phenotype-based zebrafish embryo screens led to the discovery of other classes of small molecule inhibitors acting in the BMP-pathway (Cheng et al., 2019; Dasgupta et al., 2017; Gebruers et al., 2013; Sanvitale et al., 2013).

Previously, we performed a large-scale screen of over 10,000 fungal filtrates on developing zebrafish embryos (Hoeksma et al., 2019). Embryos treated with fungal filtrate were compared to an untreated control and filtrates were scored as positive if any developmental defects were observed at 48 hours post fertilization (hpf). Here, we describe identification of cercosporamide from one of these fungi, which induced developmental defects reminiscent of BMP inhibition. We demonstrate that cercosporamide inhibited BMP/SMAD signaling in cells and zebrafish embryos. Moreover. Using kinase assays with purified kinases, cercosporamide was found to be a direct inhibitor of BMP type I receptor kinase activity. Our results indicate that cercosporamide is a bona fide BMP type I receptor inhibitor in zebrafish and mammalian cultured cells.

## Results

### Purification and identification of cercosporamide

In a screen of over 10,000 fungal filtrates on developing zebrafish embryos, we found that the filtrate of *Ascochyta aquilegiae* (CBS 168.70) induced characteristic defects, including lack of the ventral fin at the posterior part of the tail at 48 hours post fertilization (hpf). In addition, several embryos displayed the formation of secondary tissue, compared to non-treated control (Fig. 1A,B). This phenotype is strikingly similar to the phenotype induced by known BMP type I receptor kinase inhibitors such as dorsomorphin, LDN-193189 and DMH-1 (Fig. 1C), and to BMP mutants as previously reported in multiple studies (Gebruers et al., 2013; Yang and Thorpe, 2011; Yu et al., 2008).

**Fig. 1.**
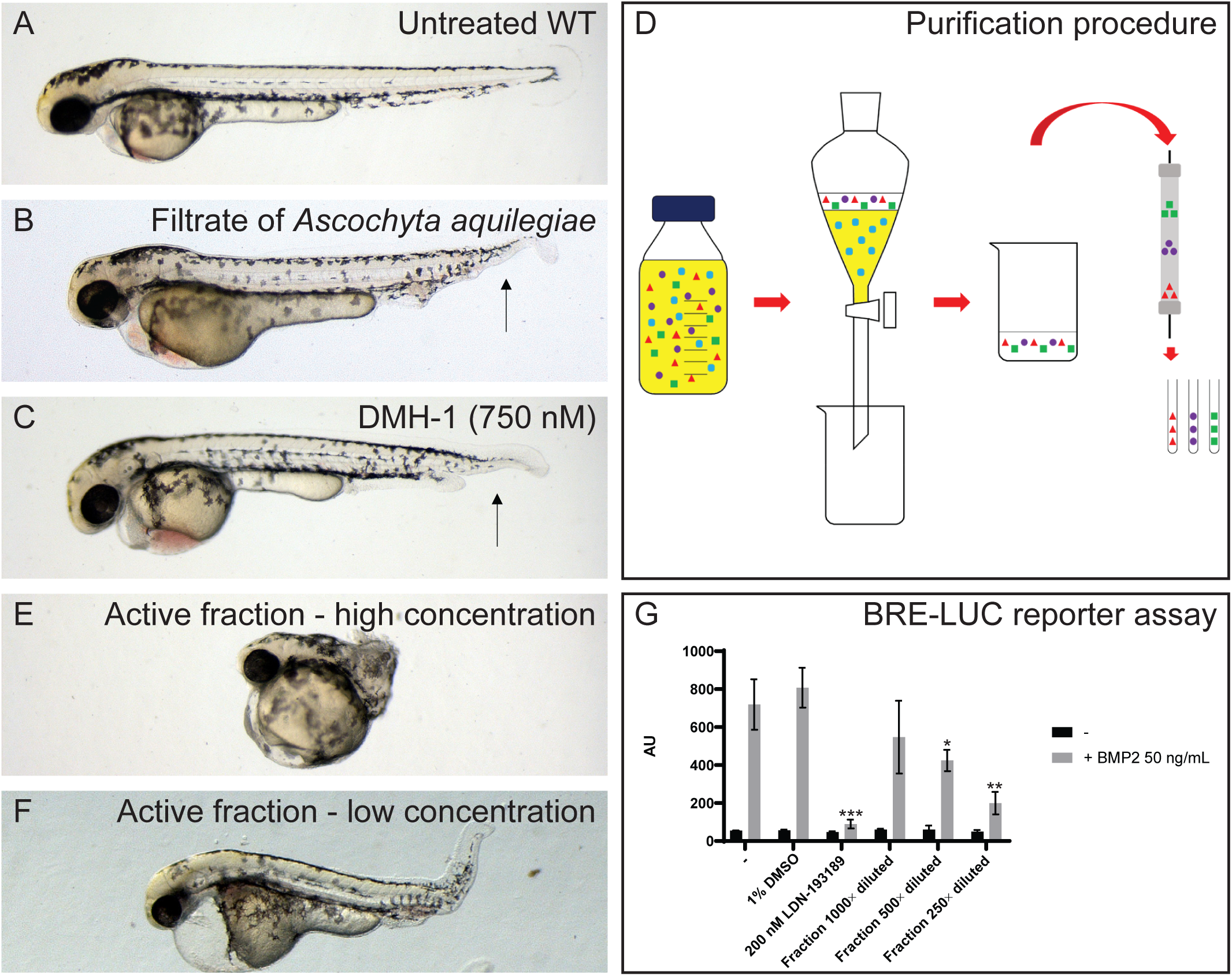
Identification of a fungal filtrate with activity consistent with inhibition of BMP signaling. (**A**) Untreated control, not treated with fungal filtrate. (**B**) An example of a phenotype induced by the filtrate of *A. aquilegiae* mixed in 1:1 ratio with E3-medium, treatment 6-48 hpf. (**C**) Phenotype induced by 750 nM DMH-1. The arrows in both B and C are indicating the loss of the ventral fin. (**D**) Schematic overview of purification of active component. The filtrate is extracted with 3× 1/3 volume ethylacetate. The ethylacetate fractions are than combined and dried. The residue is dissolved in DMSO and subsequently brought on to a preparative HPLC column. Fraction are collected every 63 seconds. (**E**,**F**) Phenotypes induced by purified fraction, high and low concentrations respectively. (**H**) Dose dependent inhibitory effect of purified fraction of fungal filtrate on BMP2-induced Smad1/5-dependent BRE-luc transcriptional reporter activity. Lack of significant effect of vehicle control DMSO and potent antagonizing effect of BMP type I receptor kinase inhibitor LDN-193189 are included. Results are expressed as mean +-SD, *p<0.05, **p<0.01, ***p<0.001.

To identify the active compound in the fungal filtrate, we generated 5 L of filtrate and performed activity guided purification (Fig. 1D). First, we performed liquid-liquid extraction and tested the resulting products. We established successful extraction of the active components as the extract induced a similar phenotype as the filtrate. Next, we fractionated the extract using preparative high performance liquid chromatography (HPLC) and tested the consequent fractions in the zebrafish phenotypic assay. One fraction induced a severely truncated phenotype in zebrafish embryos, which upon dilution turned out to induce a similar phenotype as the extract (Fig. 1E,F) and hence contained the active compound(s).

Next, we tested the active fraction for effect on a BMP2-induced SMAD1/5 dependent transcriptional reporter (BRE-luc) assay in HepG2 hepatocellular carcinoma cells. We found that the compound dose dependently inhibited BMP2-induced BRE-luc activity, like the BMP receptor kinase inhibitor LDN-193189 (Figure 1G). Taken together, these results strongly suggest that the active fungal preparation inhibited BMP signaling in zebrafish embryos and human cells.

Subsequently, the active fraction was tested on an analytical HPLC for purity and the diode array detection allowed to obtain a UV-Vis spectrum with maximum absorbance at 223 nm and 257 nm and a shoulder peak at 310 nm (Fig. S1). High resolution mass spectrometry of the active compound revealed a mass of 332.0765, which suggested several options for a molecular formula. Finally, the remainder of the fraction was dried and used for nuclear magnetic resonance (NMR) spectroscopy. The resulting spectrum (Fig. S2) matched data of cercosporamide (Fig. 2), reported by Sussman et al. very closely (Sussman et al., 2004), which was also consistent with the accurate mass measurement of 332.0765. To confirm definitively that the compound we found to induce the BMP inhibitor phenotype was cercosporamide, we obtained commercially available cercosporamide and verified its activity in zebrafish embryos (Fig. 3). In addition, we established that commercially available cercosporamide eluted from the analytical HPLC column at a similar retention time as the purified compound. *Cercosporamide*: C_16_H_13_NO_7_. HRMS: found 332.0765 (M+H), calculated 332.0770 for C_16_H_14_NO_7_. NMR (400 MHz, d_6_-DMSO): 13.55 (s, 1H); 10.56 (s, 1H); 8.25 (s, 1H); 7.54 (s, 1H); 6.22 (s, 1H); 6.14 (s, 1H); 2.57 (s, 3H); 1.73 (s, 3H) (Fig. S1). UV-Vis λ_max_: 223 nm, 257 nm, 310 nm (sh).

**Fig. 2.**
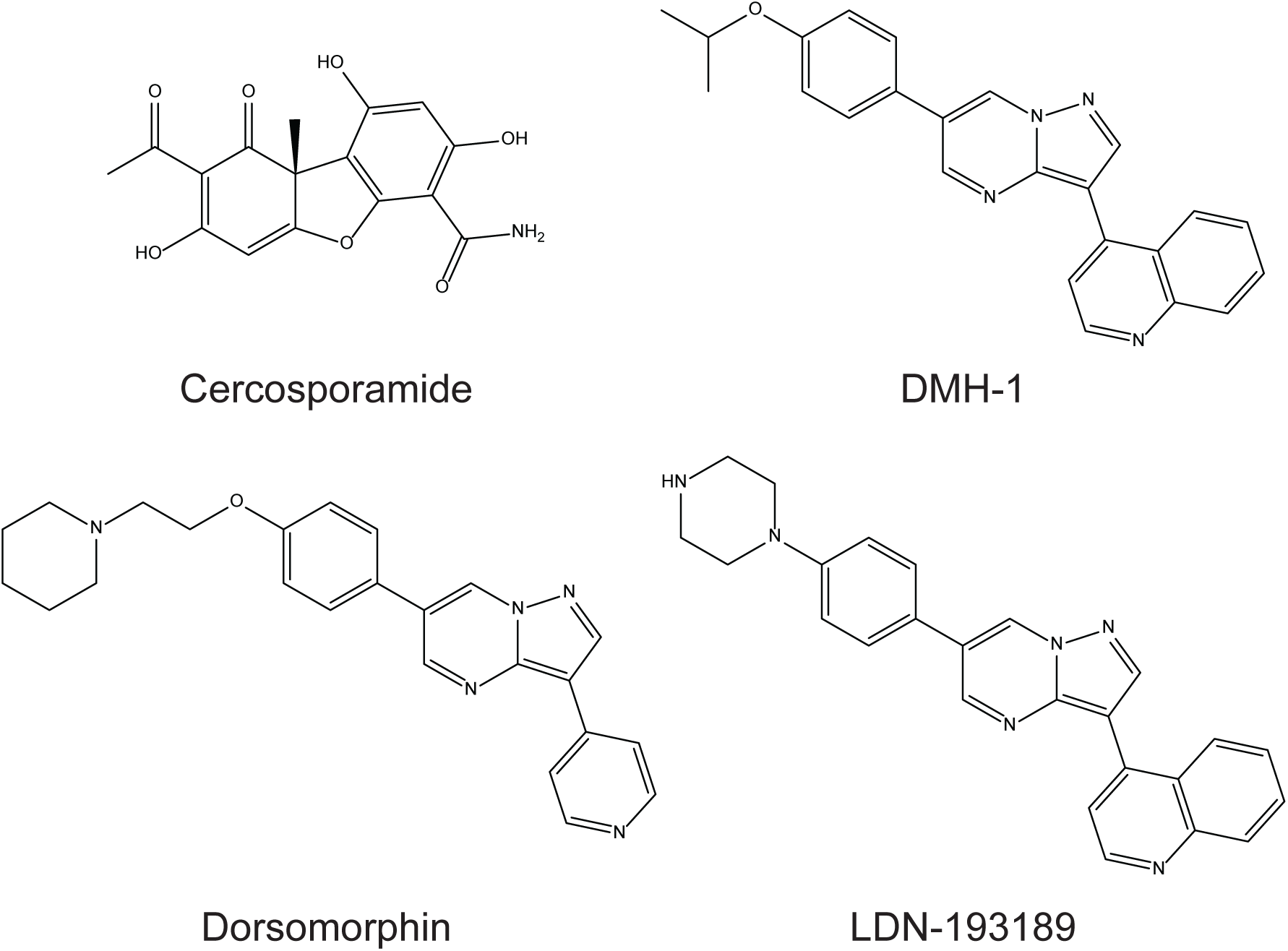
Molecular structures of cercosporamide and established BMP-inhibitors dorsomorphin, DMH-1 and LDN-193189.

**Fig. 3.**
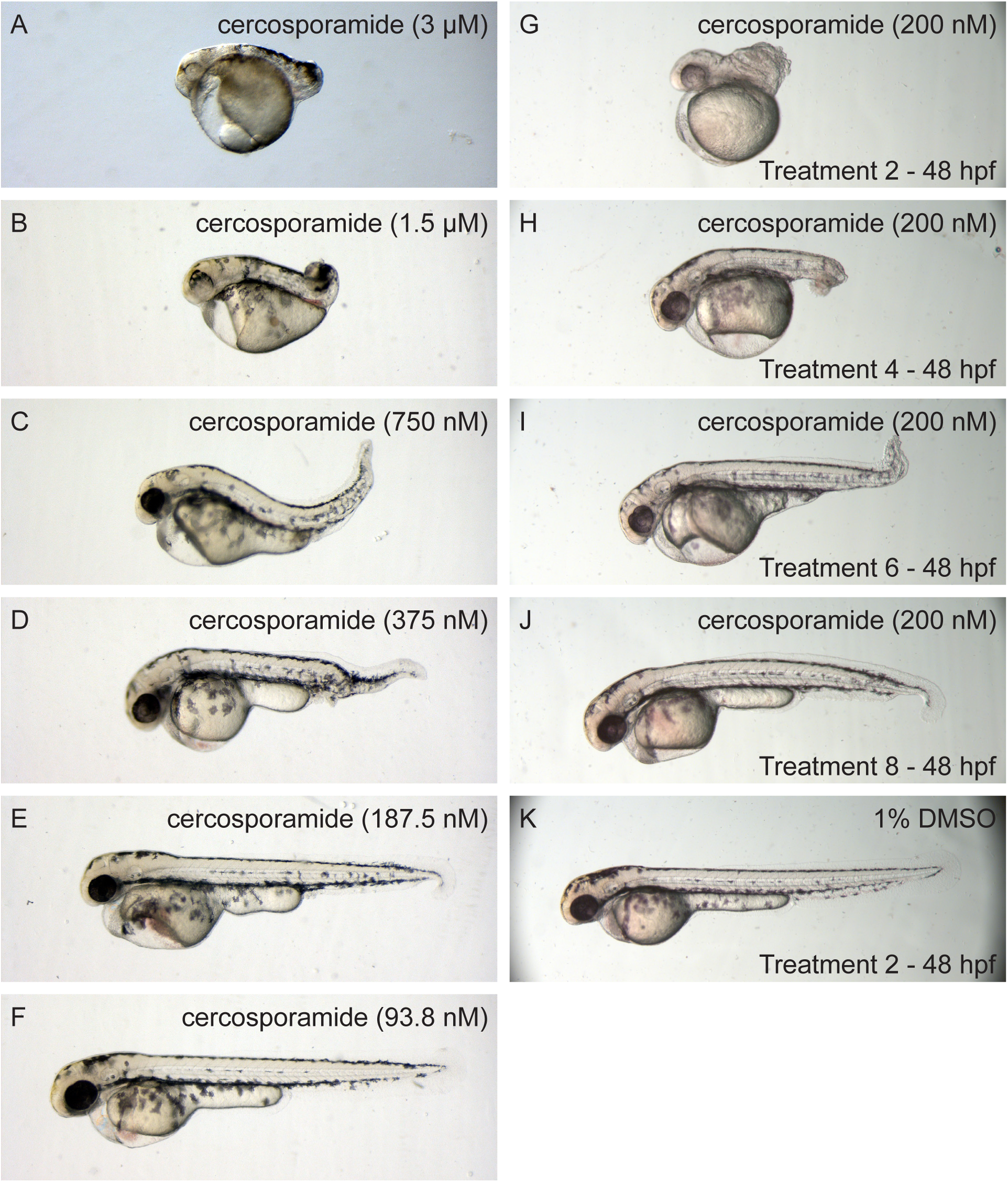
Dose- and time-dependent developmental defects of cercosporamide in zebrafish embryos. (**A-F**) Examples of phenotypes caused by a dilution range of cercosporamide (3 µM - 93 nM). (**G-J**) Examples of phenotypes caused by 200 nM cercosporamide with different treatment starting times. (**K**) Control, 1% DMSO treated embryo.

### Biological activity of cercosporamide in zebrafish assays

The biological activity of cercosporamide was tested using a dilution range in our zebrafish assay from 30.2 µM (10 µg/ml) downwards. Treatment started at 7 hpf and was continuous until effects were observed at 48 hpf. Cercosporamide treatment was lethal above 3 µM. At 3 and 1,5 µM cercosporamide induced a severe truncated phenotype (Fig. 3A,B). Similar as the purified fraction from *A. aquilegiae* (Fig. 1F), the loss-of-ventral-fin-phenotype became more evident upon dilution (Fig. 3C-E). Further dilution abolished the effect of cercosporamide at 100 nM (Fig. 3F). Furthermore, we tested the effect of starting treatment at different time points using 200 nM cercosporamide and 2 h intervals. Starting treatment at 2 hpf induced a severe truncated phenotype, comparable to treatment with 1,5 µM from 7 hpf onwards (Fig. 3G). The effect of 200 nM cercosporamide decreased dramatically when treatment was started at later time points, until only a mild phenotype was induced when treatment started at 8 hpf (Fig. 3H-K).

The phenotypes induced by cercosporamide in zebrafish embryos were remarkably similar to known BMP type I receptor kinase inhibitors, although cercosporamide is structurally distinct and does not contain the same pyrazolo[1,5-a]pyrimidine core (Fig. 2). To compare the activity of cercosporamide to known BMP type I receptor kinase inhibitors, we performed additional experiments. First, similar to Yu et al (Yu et al., 2008), we fixed embryos treated with cercosporamide or DMH-1 at 12 hpf and performed *in situ* hybridization using *krox20-* and *myod*-specific probes, which stain rhombomeres 3 and 5 and the presomitic mesoderm, respectively (Oxtoby and Jowett, 1993; Weinberg et al., 1996). Together, these are well-established markers for convergence and extension cell movements in the developing zebrafish embryo. Both cercosporamide and DMH-1 induced lateral expression of *krox20* and a more oval shape of the embryo compared to the DMSO treated control (Fig. 4). *Myod* expression was not affected as these early stages. These results are consistent with the effects of dorsomorphin on early stage zebrafish embryos (Yu et al., 2008) and with developmental defects observed in BMP-pathway mutants (Little and Mullins, 2004).

**Fig. 4.**
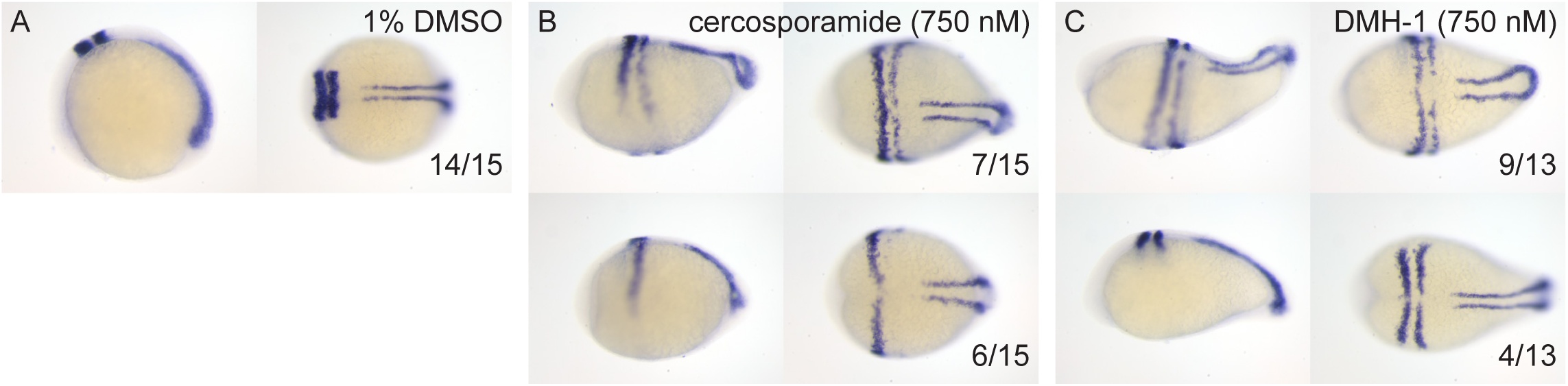
*In situ* hybridization using *krox20/myoD*-specific probes confirms that cerocsporamide and well-known BMP inhibitor, DMH-1, induced similar defects in zebrafish development. Embryos were treated with (**A**) 1% DMSO (control), (**B**) 750 nM cercosporamide, or (**C**) 750 nM DMH-1 and fixed at 12 hpf. *In situ* hybridization was done using *krox20/myoD*-specific probes. Representative examples of resulting embryos are shown with lateral view on the left and dorsal view on the right. In the bottom right corner, the fraction of embryos showing the pattern is depicted.

To assess whether cercosporamide and known BMP inhibitors exert their effects by inhibition of the same signaling pathway, we investigated whether cercosporamide and LDN-193189 cooperate by treatment of zebrafish embryos with combinations of low concentrations of cercosporamide and LDN-193189. When tested separately, 50 nM cercosporamide and 5 µM LDN-193189 did not induce detectable developmental defects. However, when tested in combination, these low concentrations induced a partial loss of the ventral fin (Fig. 5), suggesting that cercosporamide and LDN-193189 act in the same pathway. Combination treatments using different concentrations of cercosporamide and LDN-193189 always induced a more severe phenotype in the combination treatments than in the single treatments (Fig. 5). Similar results were obtained when combining cercosporamide with either dorsomorphin or DMH-1 (Fig. S3). Based on these results, we conclude that cercosporamide may exert inhibitory effects on the BMP-pathway.

**Fig. 5.**
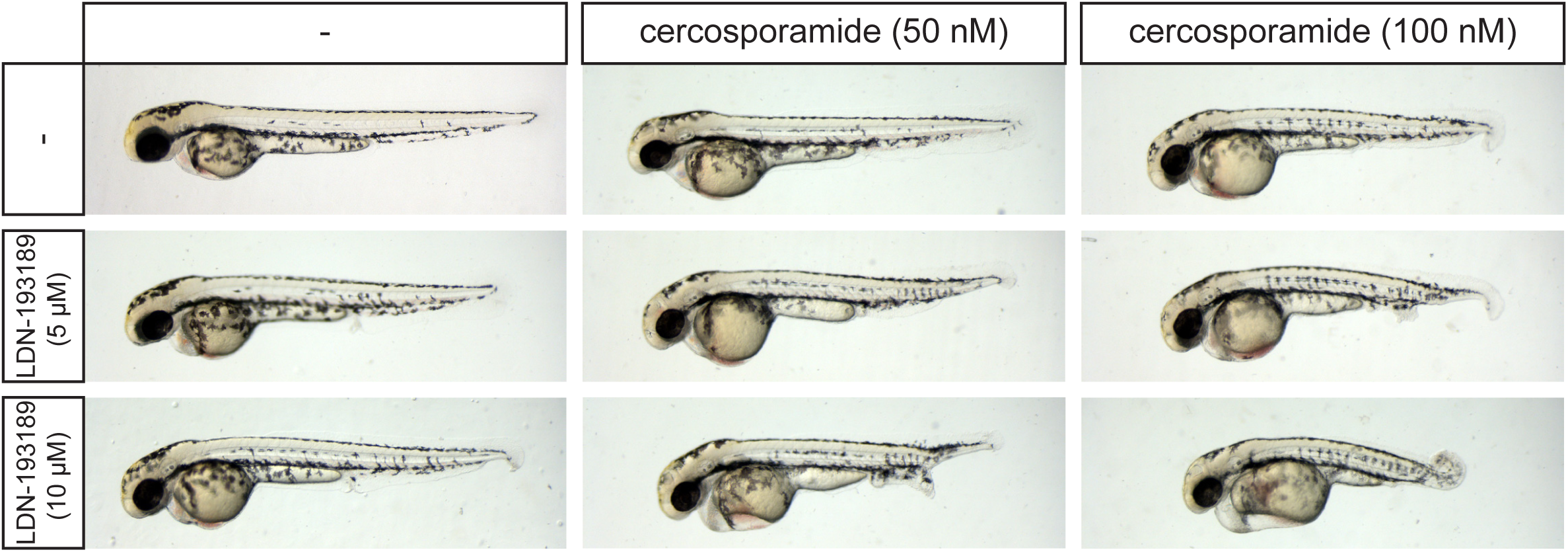
Cercosporamide and known BMP inhibitors cooperate. Combination treatments of zebrafish embryos suggest that cercosporamide acts on BMP signaling pathway. Embryos were treated with cercosporamide (50 or 100 nM) or LDN-193189 (5 or 10 µM) or combinations as indicated. Representative pictures of treated embryos are shown.

### Cercosporamide inhibits BMP-induced responses in mammalian cells

Next, we examined the effects of cercosporamide on BMP-signaling in mammalian cells. We specifically examined the inhibition of signaling through BMP2, ligand of type 1 receptors ALK3 and ALK6, and BMP6, ligand of ALK2. First, we performed a luciferase assay measuring BMP2-induced LUC expression in transfected HepG2-cells after treatment with cercosporamide (Fig. 6A). We included LDN-193189 as a positive control and DMSO as a vehicle control. Cercosporamide inhibited the BMP2-induced response in a dose dependent manner, although not as potently as LDN-193189. Subsequently, we investigated the effect of cercosporamide on BMP2-induced SMAD1/5 phosphorylation in HepG2-cells. Comparable to the luciferase assay, high concentrations of cercosporamide are capable of blocking SMAD1/5 phosphorylation, however not to same extent as 200 nM LDN-193189 (Fig. 6B). Finally, we also assessed the effect of cercosporamide on BMP6-induced BRE-reporter activity and SMAD1/5 phosphorylation. Surprisingly, we only observed minor inhibition of SMAD1/5 phosphorylation at the higher cercosporamide concentrations (Fig. 6C,D). These results demonstrate that cercosporamide inhibited BMPR-induced signaling in mammalian cells.

**Fig. 6.**
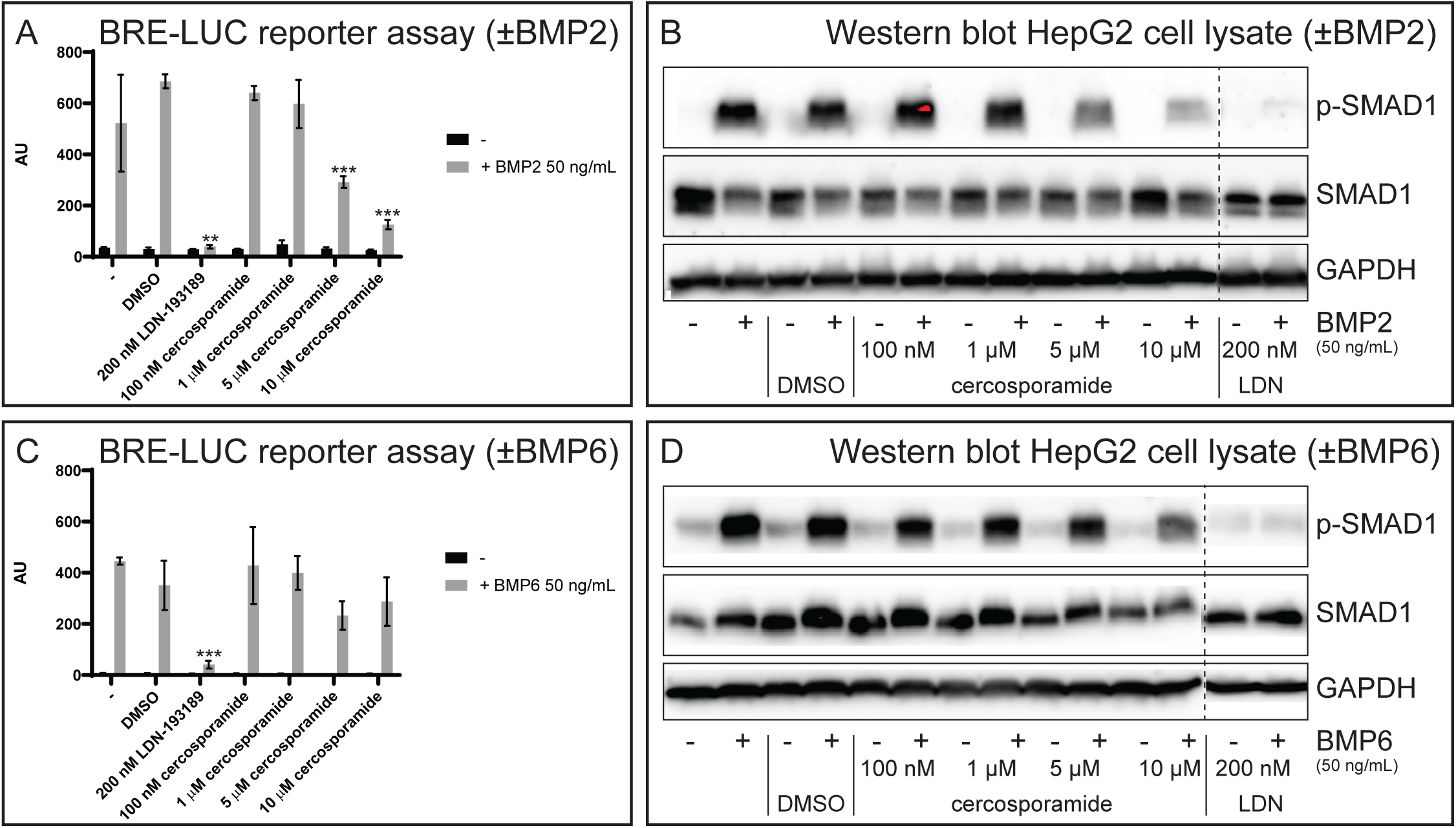
Cercosporamide inhibits BMP/Smad signaling in mammalian cells. (A) HepG2 cells with BRE-luc reporter were treated with BMP2 (50 ng/ml) or not treated (-). Control (1% DMSO), LDN-193189 (200 nM) or a range of concentrations of cercosporamide (100 nM - 10 µM) was added and luciferase activity was determined. Averages of triplicate measurements are depicted as arbitrary units. (B) HepG2 cells were treated with BMP2 (50 ng/ml) or not (-), and with 1% DMSO (control), a range of concentrations of cercosporamide (100 nM - 10 µM) or LDN-193189 (200 nM). Cells were lysed, the lysates run on SDS-PAGE gels. The material on the gel was transferred to blots and parallel blots were probed using antibodies, specific for phosphoSMAD1/5/8 (p-SMAD1) and SMAD1 (top of the blot) or GAPDH (loading control, bottom of the blot). Detection was done by enhanced chemiluminescence (ECL). Representative blots are shown. Lanes from separate parallel blots are indicated with a dashed line. (C) as in (A), except BMP6 (50 ng/ml) was used instead of BMP2. (D), as in (B), except BMP6 (50 ng/ml) was used instead of BMP2. *, ** and *** in (A) and (C) indicated p< 0.05, <0.01 and <0.001 respectively. All samples in (B) and (D) were run on same gel. Dotted line indicates where blot was cut.

### Cercosporamide is a direct BMP receptor type I kinase inhibitor

Cercosporamide may inhibit BMP signaling by inhibition of ligand-receptor interaction, by inhibiting receptor activity or by inhibiting downstream SMAD phosphorylation. Cercosporamide is reported to have serine/threonine kinase inhibitor activity with strong inhibitor activity on the cytoplasmically localized kinases, MNK1 and MNK2 (Altman et al., 2018; Konicek et al., 2011; Sussman et al., 2004). To assess whether BMPRIs were inhibited by cercosporamide, a radiometric protein kinase activity assay was performed using ALK1-ALK6. All ALKs, except for ALK1, showed an IC50 in the nano-molar range, indicating that cercosporamide might indeed act through direct inhibition of ALKs. As a control, inhibition of MNK1 and MNK2 was assessed. Indeed, MNK1 and MNK2 were strongly inhibited by cercosporamide with IC50s of 16 and 6,5 nM, respectively (Fig. 7). As a negative control, we included BMP-signaling-unrelated kinases MEK and a tyrosine kinase, anaplastic lymphoma kinase (ALK), which were indeed 7 - 67-fold less sensitive to cercosporamide than the ALK Ser/Thr kinases (Fig. 7).

**Fig. 7.**
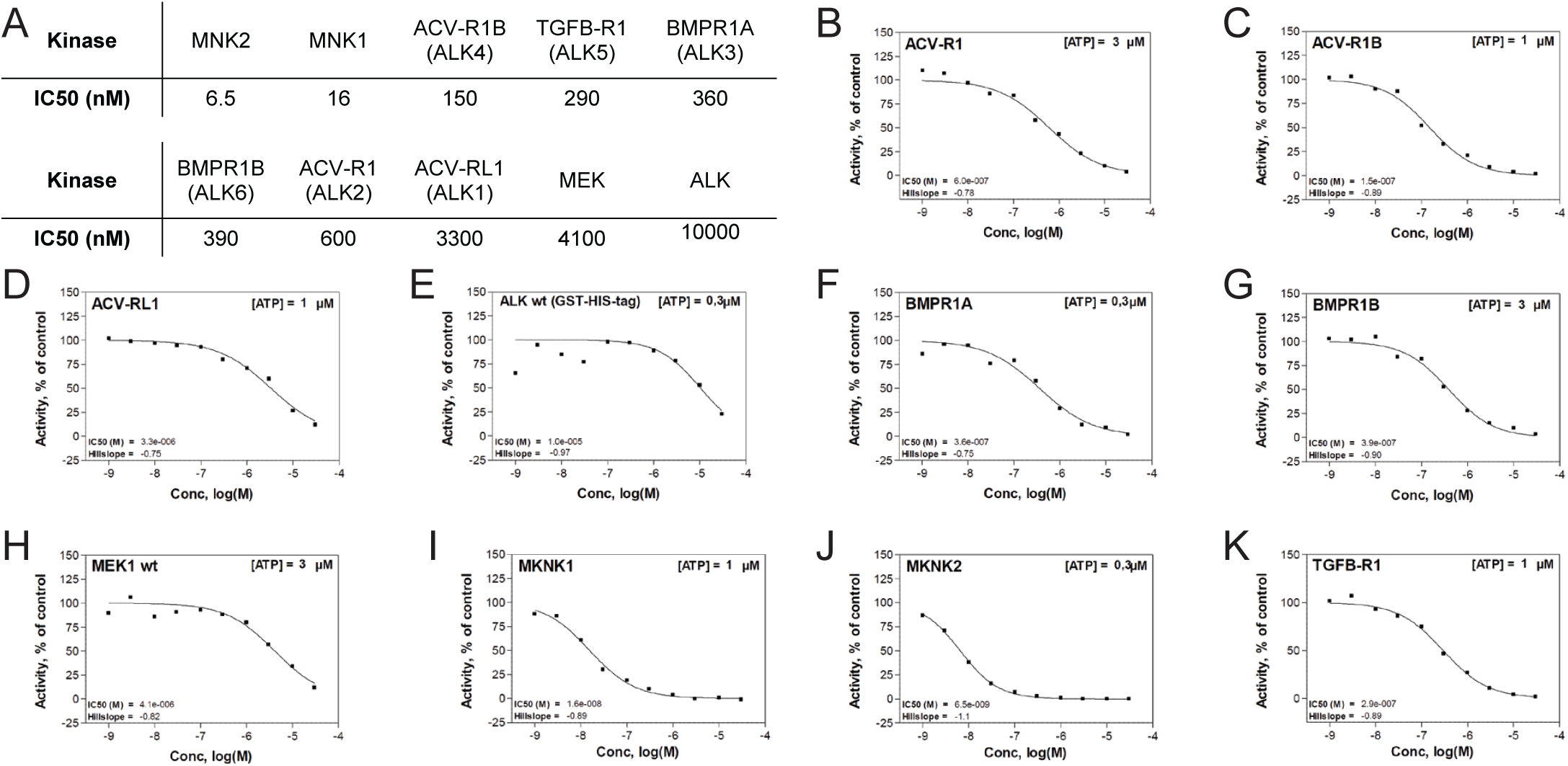
Cercosporamide inhibits kinase activity of purified ALK receptors in vitro. (**A**) IC50 values of cercosporamide-mediated inhibition of a panel of 10 kinases, as derived from the acivity graphs in panels **B-K**. The kinases include 6 type I ALK BMP receptors, mitogen-activated protein kinase (MAPK) interacting protein kinases 1 and 2 (MNK)1, MNK2, Mitogen-activated protein kinase kinase (MEK) and the tyrosine kinase, anaplastic lymphoma kinase (ALK).

Given the strong inhibitory activity of cercosporamide on MNK1 and MNK2, we wondered whether the effects of cercosporamide might be due to inhibition of MNK1 and/or MNK2. To test this, we investigated the effect of another MNK inhibitor, eFT508 on zebrafish and cultured HepG2 cells. No effect was observed at the maximum tested concentration of 20 µM on zebrafish embryo development or on SMAD1/5 phosphorylation in HepG2 cells (Fig. S4), suggesting that inhibition of MNK1 and/or MNK2 was not involved in the observed effects of cercosporamide. Together, our data are consistent with cercosporamide affecting zebrafish embryo development and mammalian cultured cells by direct inhibition of BMPRI kinase activity.

### Cercosporamide inhibits caALK signaling in mammalian cells and rescues caAlk2-induced developmental defects in zebrafish embryos

Constitutively active (ca) ALKs have been generated that signal in a ligand-independent manner. To investigate the effect of cercosporamide on different BMP type I receptors in living cells, we ectopically expressed caALK1, caALK2, caALK3 or caALK6 in HEK 293 T cells. All ca BMPRIs induced SMAD1/5 phosphorylation, albeit to different extents, which was strongly reduced by treatment with BMP inhibitor, LDN-193189 (Fig. 8A). Cercosporamide inhibited SMAD1/5 phosphorylation in response to each of these caALKs in a dose-dependent manner (Fig. 8A). We also investigated the effect of cercosporamide on caALK5, constitutively active transforming growth factor-β (TGF-β) type I receptor. Surprisingly, caALK5 induced SMAD1/5 phosphorylation was only weakly inhibited by cercosporamide (Fig. S5), even though ALK5 kinase activity was potently inhibited by cercosporamide in vitro (Fig. 7). To investigate the role of ALK5 in the effects of cercosporamide further, we tested the effect of two selective ALK5-kinase inhibitors, i.e. ALK5 kinase inhibitor II and A83-01, on zebrafish embryos. Surprisingly, both these compounds induced a severely curved tail and fused eyes in a broad concentration range (Fig S6). These developmental defects are distinct from the phenotype induced by cercosporamide, suggesting that cercosporamide does not act *in vivo* through inhibition of Alk5. Thus, cercosporamide selectively inhibits caBMP type I receptors, but not ca TGF-β-like receptors in cultured cells.

**Fig. 8.**
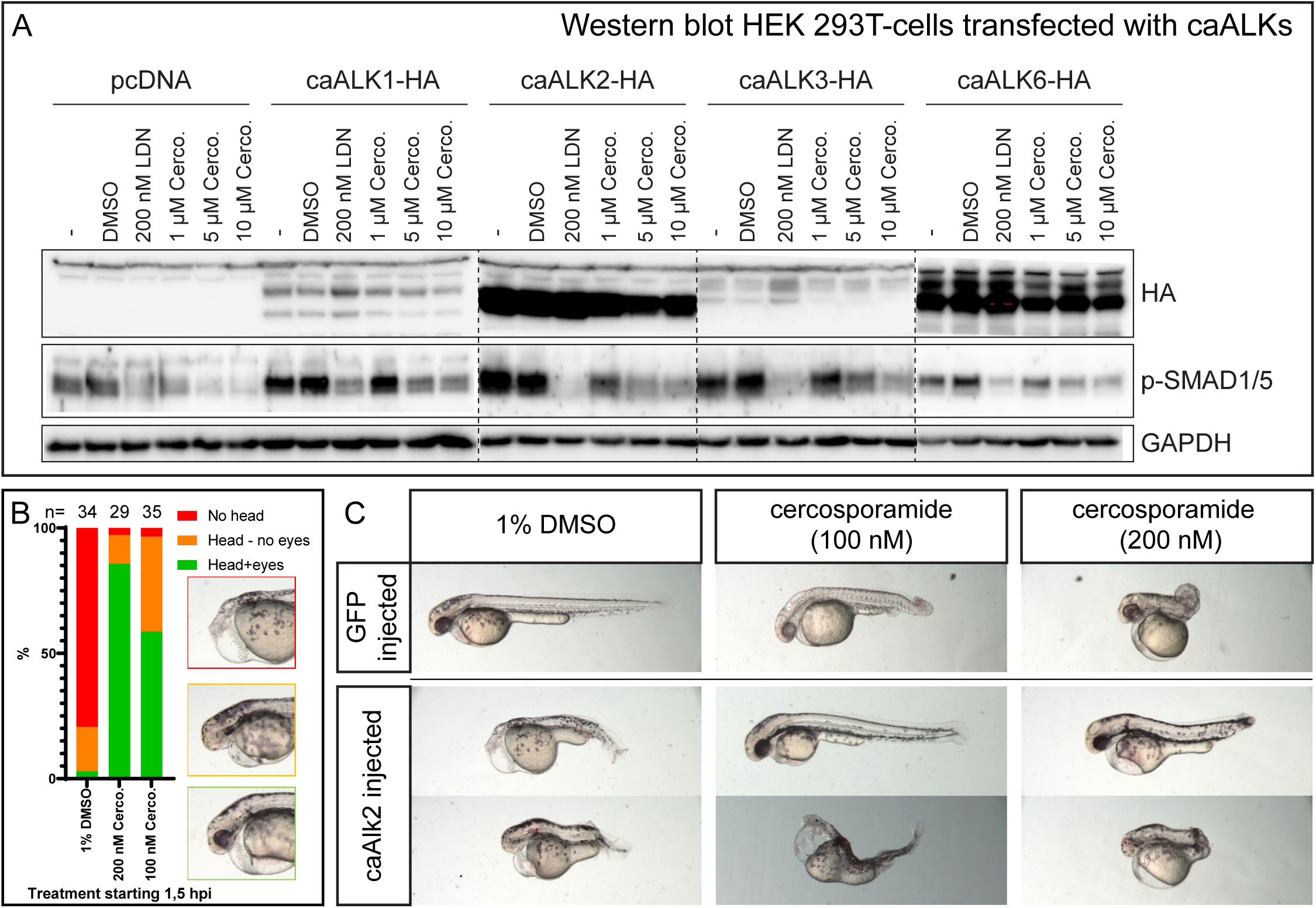
Cercosporamide selectively inhibits caBMP type I receptor in cultured mammalian cells and zebrafish embryos. (**A**) HEK 293 T cells were transfected with expression vectors for caALK1, caALK2, caALK3 or caALK6 each with a haemagglutinin (HA) epitope tag at the carboxy terminus. The cells were treated with vehicle (1% DMSO), LDN193189 (200 nM) or a range of cercosporamide concentrations (1-10 µM as indicated). Subsequently, the cells were lysed and the lysates run on SDS PAGE gels. The material on the gel was transferred to blots and parallel blots were probed with antibodies specific for HA (epitope tag on receptors), phosphoSMAD1/5/8 (p-SMAD1, top of the blot) or GAPDH (loading control, bottom of the blot). Dotted line indicates borders of different gels. (**B**) Cercosporamide partially rescued caAlk2 induced developmental defects in zebrafish embryos in vivo. Bar chart showing phenotype distribution of embryos injected with caAlk2 mRNA and subsequently treated with either 1% DMSO, 100 nM or 200 nM cercosporamide. The severity of the phenotype (examples depicted in the insets) is plotted as red (severe, no head), orange (intermediate, head structures present, no eyes) and green (mild, head structure with eyes detectable). The total number of embryos (n) is indicated. (**C**) The phenotypes of the rescued embryos are highly variable and therefore, we depicted two representative indiciduals for each condition.

Next, we examined the ability of cercosporamide to inhibit Alk2 *in vivo* in zebrafish embryos expressing caAlk2. Messenger RNA encoding caAlk2, combined with GFP mRNA was injected at the one-cell-stage. The embryos were incubated with either 100 nM or 200 nM cercosporamide or 1% DMSO (vehicle control) from 2 hpf onwards. As a control we injected GFP mRNA only. We observed a variety of phenotypes in caAlk2-injected embryos treated with DMSO, which we categorized in three groups (Fig. 8B). We observed an absence of a head in 79% of the cases when treated with DMSO (Fig 8B,C). Furthermore, of this group 18% of the embryos did develop a head, however no eyes were developed. Only 3% of the embryos develop a seemingly normal head. Treatment with cercosporamide largely rescued the head phenotype in a dose-dependent manner, in that 86% and 58% developed a head with eyes, following treatment with 200 nM and 100 nM, respectively (Fig. 8B). Conversely, the cercosporamide-induced developmental defects in the posterior region of the control GFP-injected embryos were not completely rescued by caAlk2 injection, although there appeared to be some improvement. Taken together, cercosporamide partially rescued the effects of caAlk2 injection and caAlk2 injection partially rescued the posterior defects induced by cercosporamide, indicating that cercosporamide is an inhibitor of Alk2 *in vivo*.

## Discussion

Here, we describe the purification and identification of cercosporamide from *Ascochyta aquilegiae*. The biological activity in developing zebrafish embryos suggested that cercosporamide is an inhibitor of BMP signaling. Subsequent analyses indicated that cercosporamide inhibited kinase activity of BMPRI receptor proteins *in vitro*, inhibited BMPRI signaling in mammalian cells and rescued the caALK2-induced developmental defects in zebrafish embryos *in vivo*.

Cercosporamide is a known secondary metabolite of fungi. Previously, it was purified and identified from a strain of the fungus *Cercosporidium henningsii* (Sugawara et al., 1991) and has further been found to be produced by fungal strains of the *Lachnum* and *Pseudaegerita* genus (Hosoya et al., 2011). This is the first time that cercosporamide is described as a metabolite from *A. aquilegiae*.

Initially, cercosporamide was considered to be a potent antifungal agent and phytotoxin (Sugawara et al., 1991). Later, it was shown that cercosporamide inhibits PKC1 in yeast (Sussman et al., 2004). Furthermore, cercosporamide has been tested on a panel of kinases and was identified as a potent MNK1 and MNK2 inhibitor. Inhibition of MNK1 and MNK2 kinases may be the underlying mechanism for cercosporamide-mediated suppression of growth of human hepatocellular carcinoma and acute myeloid leukemia precursors (Altman et al., 2018; Konicek et al., 2011; Liu et al., 2016). It is noteworthy that ALK4 was also included in the panel of kinases and was found to be inhibited by cercosporamide (Konicek et al., 2011), consistent with our results (Fig. 7). However, this observation has not been pursued any further and at the time, no functional assays were done to test the hypothesis that cercosporamide exerts its function through inhibition of TGF-β family type I receptor kinases.

Using the zebrafish embryo model, we identified cercosporamide as a potent BMP-inhibitor. Cercosporamide mimics the phenotype induced by established BMP-inhibitors and the phenotype observed in zebrafish mutants with loss-of-function mutations in factors of the BMP signaling pathway. Moreover, we showed that cercosporamide rescued developmental defects induced by caALK2 overexpression in a similar fashion as DMH-1. The molecular structure of cercosporamide is completely different than the structure of established BMP-inhibitors, which mostly contain a pyrazolo[1,5-a]pyrimidine core. Hence, our results with cercosporamide may unlock an entirely different class of molecules that may be used as BMP-inhibitors, potentially through a distinct working mechanism. Future follow-up analyses to elucidate the interaction of cercosporamide with its targets at the structural and molecular level will provide insight into the mode of action of cercosporamide. Our work clearly underlines the value of performing small molecule compound screens on zebrafish embryos in order to uncover potential new drugs.

Our data are consistent with cercosporamide acting through inhibition of ALK2. Yet, ALK2 was not the most potently inhibited BMP receptor in the *in vitro* kinase assays (Fig. 6). ALK4 and ALK5 were inhibited more efficiently, consistent with published data (Konicek et al., 2011). However, cercosporamide induced different developmental defects than renowned Alk5-inhibitors in zebrafish (Fig. S5). In the reporter assays, ALK3 and ALK6 appeared to be more potently inhibited than ALK2. On the other hand, the developmental defects induced by cercosporamide in zebrafish embryos have a striking similarity to zebrafish mutants that lack functional Alk2 (Bauer et al., 2001; Mintzer et al., 2001). Moreover, caAlk2-induced developmental defects were rescued by cercosporamide in zebrafish embryos *in vivo*. It appears that the *in vivo* effects of cercosporamide not only depend on the specificity for distinct ALKs, but also on other factors that determine the function of the different ALKs in development. *In vivo*, cercosporamide may exert its effects mainly through inhibition of Alk2.

Taken together, our results suggest that cercosporamide and possibly derivatives of cercosporamide have the potential to be used as BMP-inhibitors, thus unlocking a new class of molecules that may be developed further for use in a clinical setting, for instance to combat diseases with overactive BMPR signaling, including FOP and DIPG.

## Materials and Methods

### Zebrafish embryo assay

Zebrafish eggs obtained from family crosses of Tuebingen Long fin zebrafish lines were used to assess the biological activity of all samples. The eggs were washed with fresh E3-medium and subsequently divided over 24-well plates, 10 embryos per well in 1000 µL E3-medium. Samples were added to the wells at various times as mentioned in text and Figures. At 48 hours post fertilization (hpf), the zebrafish embryos were inspected for morphological developmental defects. Embryos displaying morphological defects were imaged using a Leica MZFLIII microscope equipped with a Leica DFC320 camera or Leica M165 FC microscope equipped with a Leica DMC5400 camera.

All procedures involving experimental animals were approved by the local animal experiments committee (Koninklijke Nederlandse Akademie van Wetenschappen-Dierexperimenten commissie) and performed according to local guidelines and policies in compliance with national and European law. Adult zebrafish were maintained as previously described (Aleström et al., 2019).

### Culture and isolation of active compound

Initially, the fungus *Ascochyta aquilegiae* (CBS 168.70) was grown on a cornmeal agar plate for 7 days at 25 °C. The plate with mycelium was then cut into cubes of approximately 5×5 mm. Subsequently, a bottle of 100 mL containing 50 mL Czapek Dox Broth+0.5% yeast extract was inoculated with two cubes and incubated at room temperature (RT) for 7 days. The medium was filter sterilized using a 0.45 µm Millipore filter and tested in serial dilution in the zebrafish embryo assay. In order to increase the yield of the active components, we optimized the growth conditions for this fungus before generating a large batch of filtrate. Ultimately, we inoculated 100 bottles containing 50 mL Czapek Dox Broth without the addition of yeast extract and incubated the medium at 15 °C for 14 days. The medium was filtered as mentioned above in batches of 1 L each.

Subsequently, each liter was extracted with 3x ±300 ml ethyl acetate (EtOAc). The EtOAc was combined and evaporated using a rotation evaporator. The residue was dissolved in 2 ml dimethylsulfoxide (DMSO), of which a small aliquot was used in the zebrafish embryo assay to verify the successful extraction of the active components. Successively, the extract was fractionated on a modular preparative high performance liquid chromatography (HPLC) system, consisting of a Shimadzu CBM-20A controller, a Shimadzu LC-20AP pump and a Shimadzu FRC-10A fraction collector using a C18 reversed phase Reprosil column (10 µm, 120 Å, 250 × 22 mm) and a Shimadzu SPD-20A ultraviolet light (UV)-detector set at 214 nm and 254 nm. The mobile phase was 0.1% trifluoroacetic acid in acetonitrile:water 5:95 (buffer A) and 0.1% trifluoroacetic acid in acetonitrile:water 95:5 (buffer B). A flow rate of 12.5 ml min^-1^ was applied using the following protocol: 100% buffer A for 5 minutes followed by a linear gradient of buffer B (0-100%) for 40 minutes, 100% buffer B for 5 minutes, another linear gradient of buffer B (100-0%) for 5 minutes and finally 100% buffer A for 5 minutes. Fractions were collected every 63 seconds, resulting in 57 fractions of 13 ml. 1ml of each collected fraction was dried in a speed-vac overnight. The fraction residues were dissolved in 50 µL in DMSO and tested in serial dilutions starting at 100× diluted. The sole active fraction has been analyzed using analytical chemical methods as described below.

### Identification of biologically active compounds

First, active fraction was assessed for its purity through analytical HPLC, using a Shimadzu LC-2030 system with Photodiode Array (PDA) detection (190-800 nm) using a Shimadzu Shim-pack GIST C18-HP reversed phase column (3 µm, 4.6 × 100 mm). Simultaneously, through PDA detection a UV-Vis spectrum was obtained for the active compound. Secondly, high resolution mass spectrometry (HRMS) was measured on an LCT instrument (Micromass Ltd, Manchester UK). The sample was mixed with sodium formate, allowing the sodium formate to be used as internal calibrant and facilitated identification of the more accurate mass of the compound. The remainder of the active fraction was dried in a speedvac and dissolved in 400 mL DMSO-d6. Next, a ^1^H-nuclear magnetic resonance (NMR) spectrum was measured at 400 MHz using an Agilent-400 instrument.

### Compounds

Cercosporamide, dorsomorphin, DMH-1, LDN-193189, Alk5-inhibitor II, A83-01 and DMSO were all purchased from Sigma-Aldrich (Zwijndrecht, The Netherlands). eFT508 was obtained from Toronto Research Chemicals (Toronto, Canada).

### *In situ* hybridization

Embryos were treated from approximately 6 hpf until 12 hpf when they were fixed in 4% paraformaldehyde overnight. Furthermore, the *in situ* hybridization was performed using krox20 and myoD anti-sense RNA-probes as described elsewhere (Thisse and Thisse, 2008).

### Kinase activity assay

A radiometric protein kinase assay was performed by ProQinase GmbH (Freiburg, Germany), using purified bacterially expressed human kinases and a range of concentrations of cercosporamide. IC_50_ values were calculated using Prism 5.04 for Windows (Graphpad, San Diego, California, USA; www.graphpad.com). The mathematical model used was “Sigmoidal response (variable slope)” with parameters “top” fixed at 100% and “bottom” at 0 %. The fitting method used was a least-squares fit.

### Mammalian cell lines and treatment

HepG2 and HEK 293 T cells were routinely grown in DMEM supplemented with 10% fecal calf serum (FCS), supplemented with penicillin and streptomycin and glutamine. HepG2 and HEK 293T cells lines were obtained from ATCC, and were frequently tested for absence of mycoplasma and cell lines were authenticated using STR profiling kit from Promega.

For BMP stimulation the HepG2-cells (approximately 50% confluency) were starved on serum free medium for 6 h. Subsequently, prior to addition of BMP ligands, the cells were treated with compound for 30 minutes. Next, the cells were stimulated with BMP2 (50 ng/ml) or BMP6 (50 ng/ml) for 45 minutes. The cells were then washed with PBS and lysed in Laemmli sample buffer for Western blot analysis.

### Immunoblot Analysis

HepG2 or 293T cells were lysed in Laemmli sample buffer. Proteins were separated by sodium dodecyl sulfate polyacrylamide gel electrophoresis (SDS-PAGE) and transferred onto 45-μm polyvinylidene difluoride (PVDF) membrane (IPVH00010, Merck Millipore). Membranes were blocked using 5% non-fat dry milk in Tris-buffered saline with 0.1% Tween 20 (655204, Merck Millipore) and probed with the respective primary and secondary antibodies. The signal was detected using Clarity(tm) Western ECL Substrate (1705060, Bio-Rad) and ChemiDoc Imaging System (17001402, Bio-Rad). The antibodies used for immunoblotting were raised against the following proteins: phospho-SMAD1/5/8 (Persson et al., 1998), SMAD1 (Cell Signaling Technology), GAPDH (Merck Millipore) and HA (12CA5; Roche).

### Transfections, luciferase assays and DNA constructs

For luciferase transcriptional reporter assays, HepG2 cells were seeded in 9 cm plates at approximately 60% confluency and transfected with polyethyleneimine (PEI). Twenty-four hours later the transfected cells were seeded in 24 wells plates at approximately 60% confluency. Another 24 hours later the cells were serum starved for 6 hours. Subsequently the cells were treated with compound or DMSO for 30 minutes. Thereafter the cells were stimulated with BMP2 in the presence of compounds (or DMSO) for 16 hours (o/n). Subsequently the cells were washed with PBS and lysed. Luciferase activity was measured using the luciferase reporter assay system from Promega (Leiden, The Netherlands) by a Perkin Elmer luminometer Victor^3^ 1420. Each DNA transfection mixture was equalized with empty vector when necessary and every experiment was performed in triplicate. β-galactosidase expression construct was co-transfected and b-galactosidase was measured to normalize for differences in transfection efficiency. The BRE-Luc reporter construct has been reported before (Korchynskyi and Ten Dijke, 2002).

For experiments with the constitutively active (ca) ALK type 1 receptors constructs, HEK 293 T cells were seeded in 24 wells at approximately 90% confluency and transfected with the DNA expression constructs in the presence of polyethylenimine (PEI). Thirty hours after transfection the cells were put on serum starved medium and treated with compounds for 16 hours (o/n). Subsequently the cells were washed with PBS and lysed in Laemmli sample buffer. Expression constructs for constitutively active (ca) type I receptors (caALK1, caALK2, caALK3, caALK5, caALK6) were previously described (Dennler et al., 1998; Fujii et al., 1999).

### mRNA synthesis and micro-injection

The pCS2+ plasmid encoding constitutively active Alk2 with a Glutamine to Aspartic acid substitution at position 204 was kindly donated by Jeroen Bakkers (Smith et al., 2009). The DNA sequence of the inserts in plasmid constructs were verified. Both plasmids were digested with *Not*I and mRNA was generated with SP6 RNA polymerase using mMessage mMachine kit (Ambion). The mRNA was purified through phenol/isoamylalcohol/chloroform extraction. Zebrafish embryos were injected into the yolk at the one cell stage with approximately 1 nL of either 50 ng/µL green fluorescent protein (GFP) mRNA or a cocktail containing 10 ng/µL caAlk2 and 50 ng/µL GFP mRNA. Subsequently, the embryos were washed with E3-medium and distributed in a 12-well plate, 15-20 embryos per well, and incubated with either DMSO, 100 or 200 nM cercosporamide from 2 hpf onwards. Next, embryos were selected for fluorescence at 24 hpf and examined at 48 hpf. The phenotypes were categorized in three groups: no head; head, no eyes, head and eyes. Finally, the bar graph was generated using Prism 8.3.0 for Windows (GraphPad, San Diego, CA, USA). The represented data is a combination of two repeats of the experiment.

## Acknowledgements

We like to thank Jeroen Bakkers for zebrafish caAlk2 cDNA construct. We like to thank Albert Heck and Arjan Barendregt for their help with HRMS measurements and Geert-Jan Boons and Justyna Dobruchowska for their help with NMR measurements. This study was supported by Cancer Genomics Centre Netherlands (CGC.NL to PtD).

## Author contributions

Conceived and designed the experiments: JH, GvdZ, PtD, JdH. Performed the experiments: JH, GvdZ. Analyzed the data: JH, GvdZ, PtD, JdH. Wrote the paper: JH, PtD, JdH.

## Supplementary figures

**Fig. S1:**
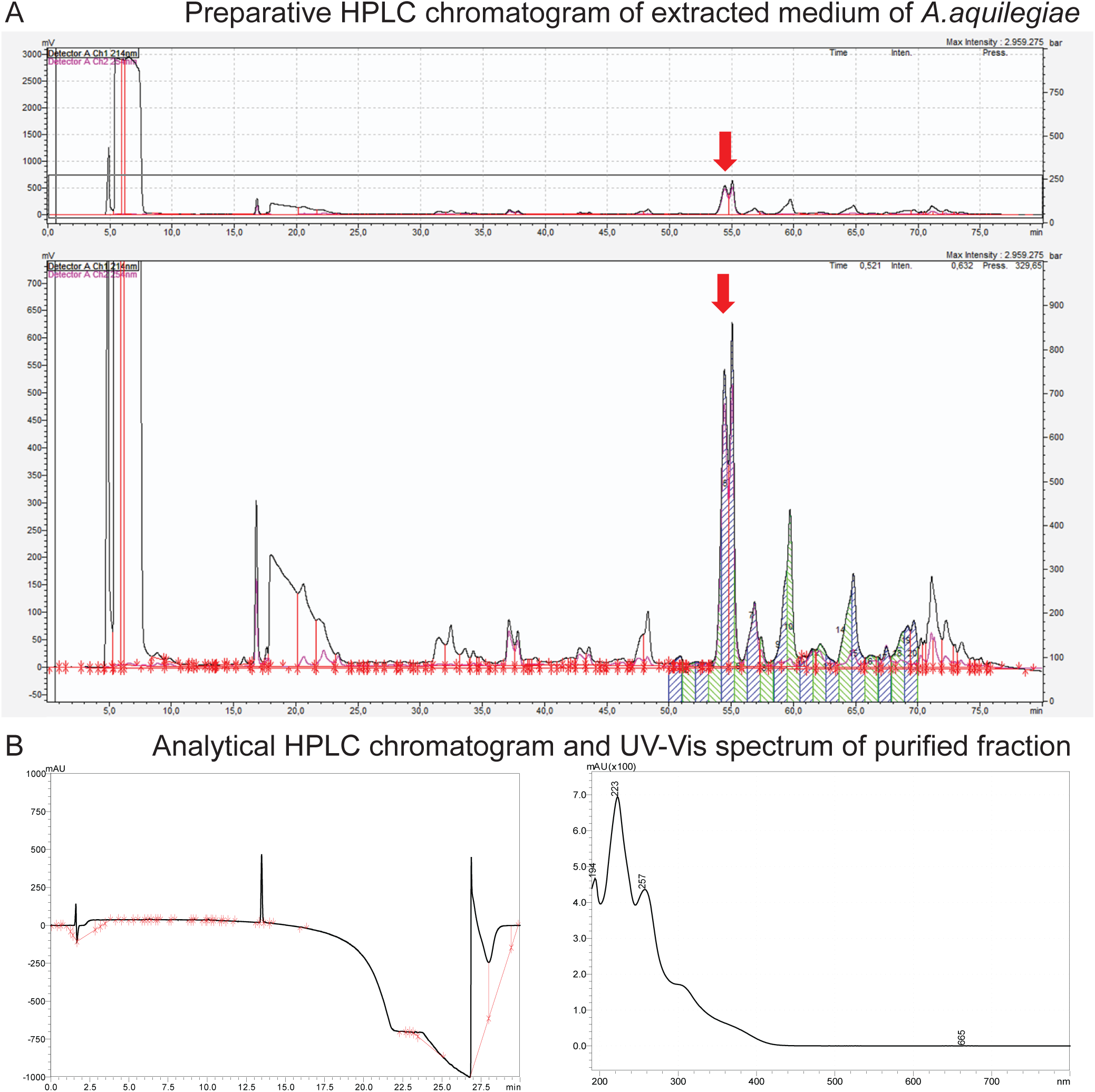
(A) Purification of active fraction (red arrow) through preparative HPLC. (B) Assessment of the purity and aquirement of an UV-Vis spectrum of the active fraction through analytical HPLC with diode array detection.

**Figure S2:**
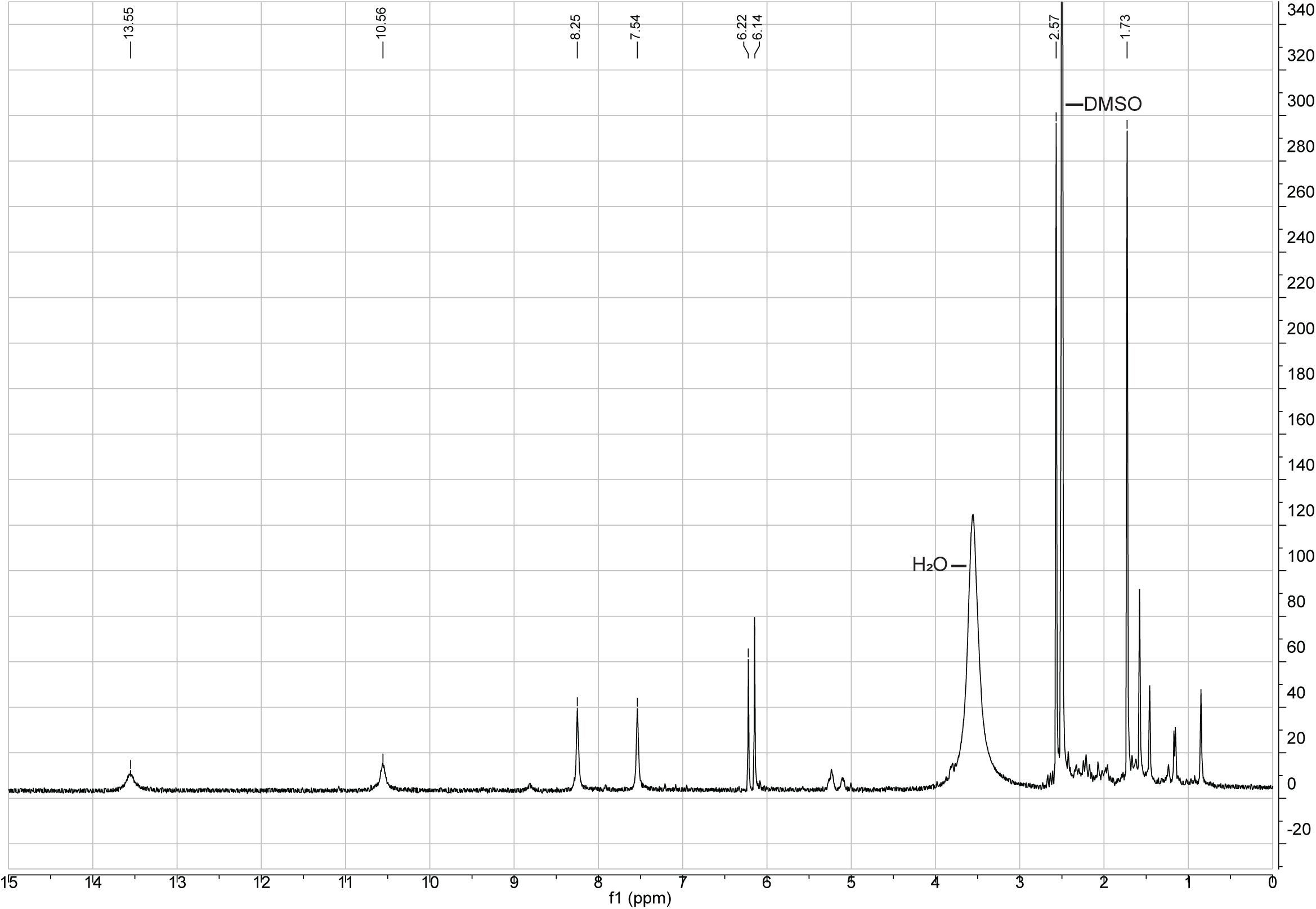
^1^H-spectrum of active preparative HPLC fraction in DMSO-d6

**Fig. S3:**
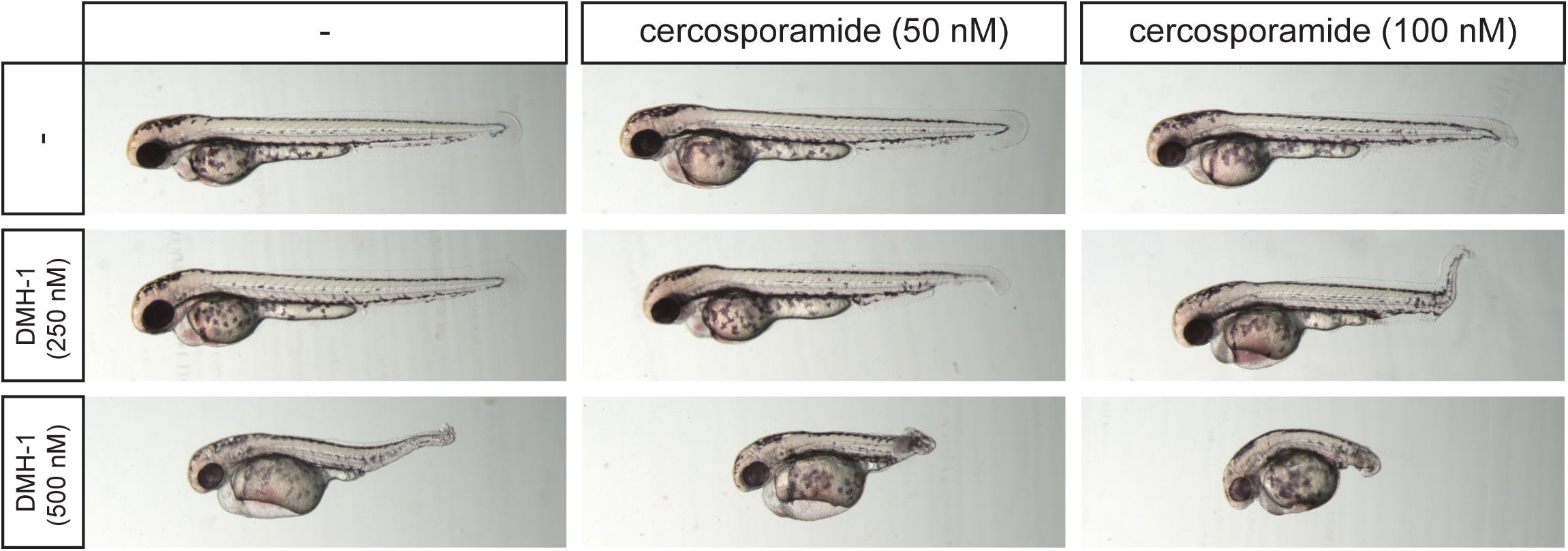
Cercosporamide and known BMP inhibitors cooperate. Combination treatments of zebrafish embryos suggest that cercosporamide acts on BMP signaling pathway. Embryos were treated with cercosporamide (50 or 100 nM) or DMH-1 (250 or 500 nM).

**Fig. S4:**
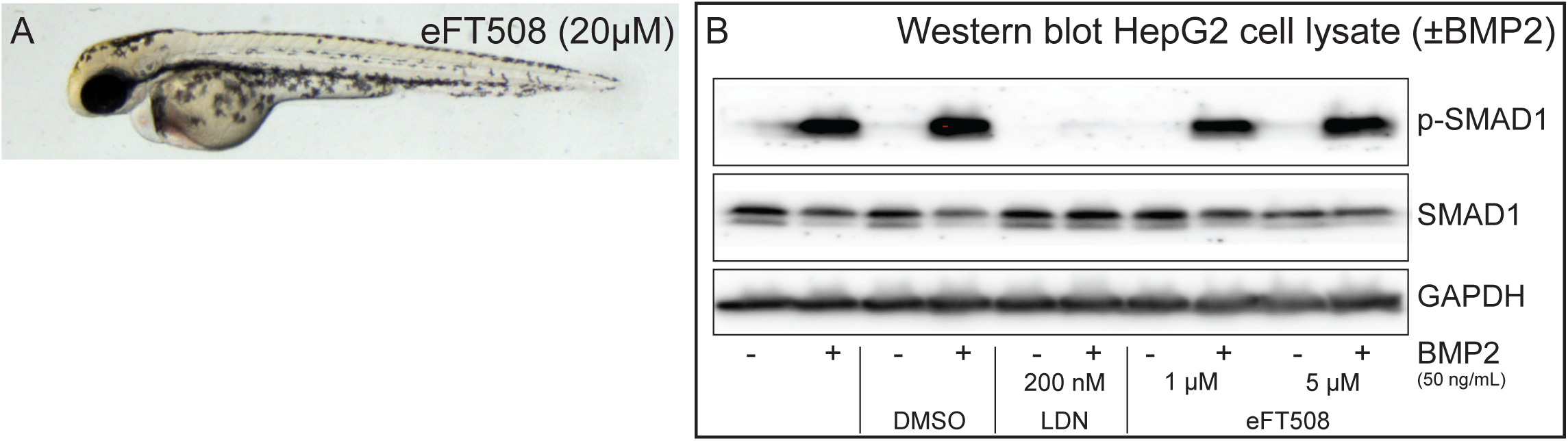
Inhibition of Mnk1/Mnk2 using the potent, selective inhibitor, eFT 508 (A) did not induce developmental defects in zebrafish embryos at 20 µM, and (B) did not affect BMP2-induced SMAD 1 phosphorylation in HepG2 cells.

**Fig. S5:**
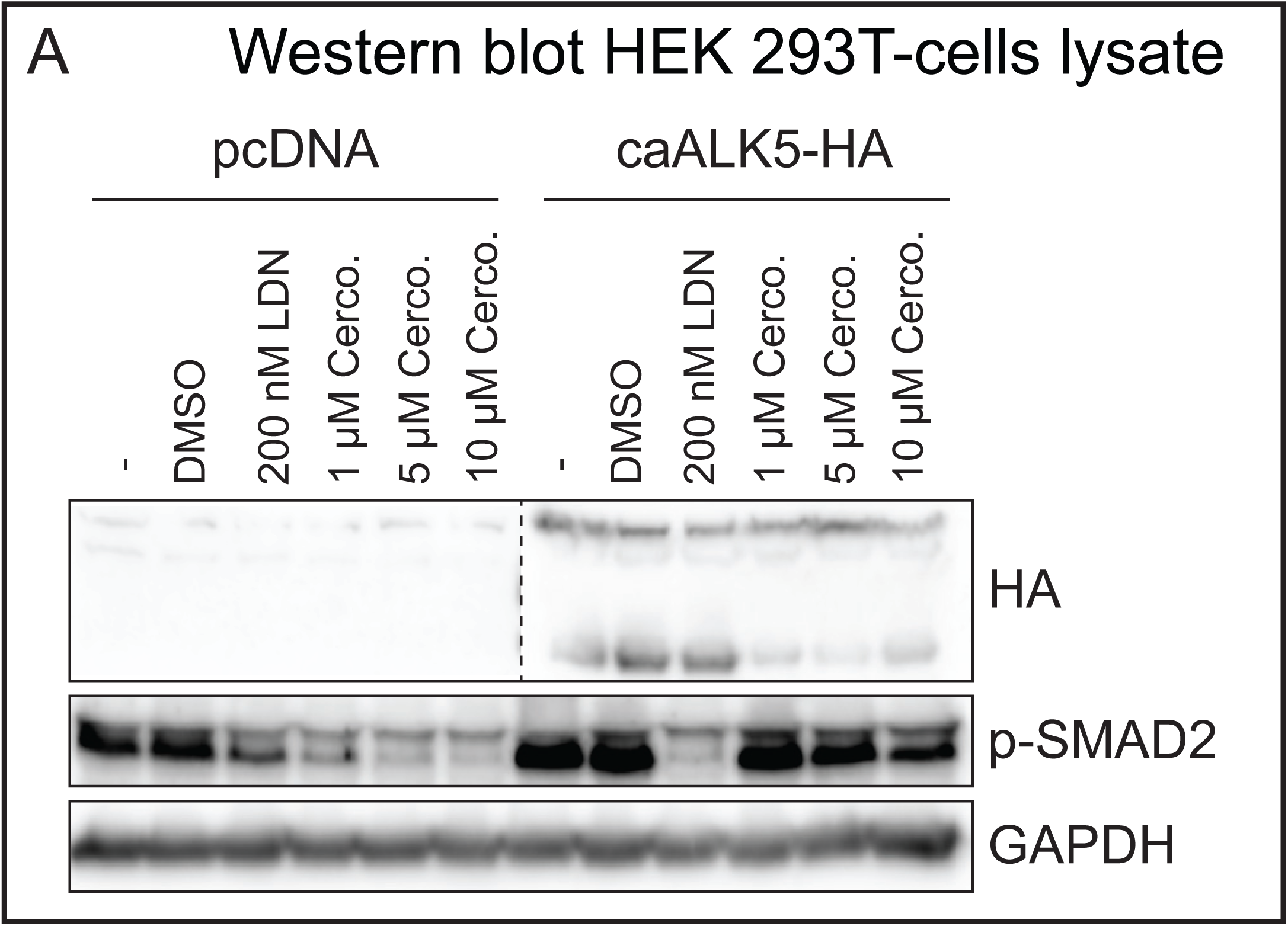
caAlk5 was not inhibited by cercosporamide in transfected HEK 293T-cells.

**Fig. S6:**
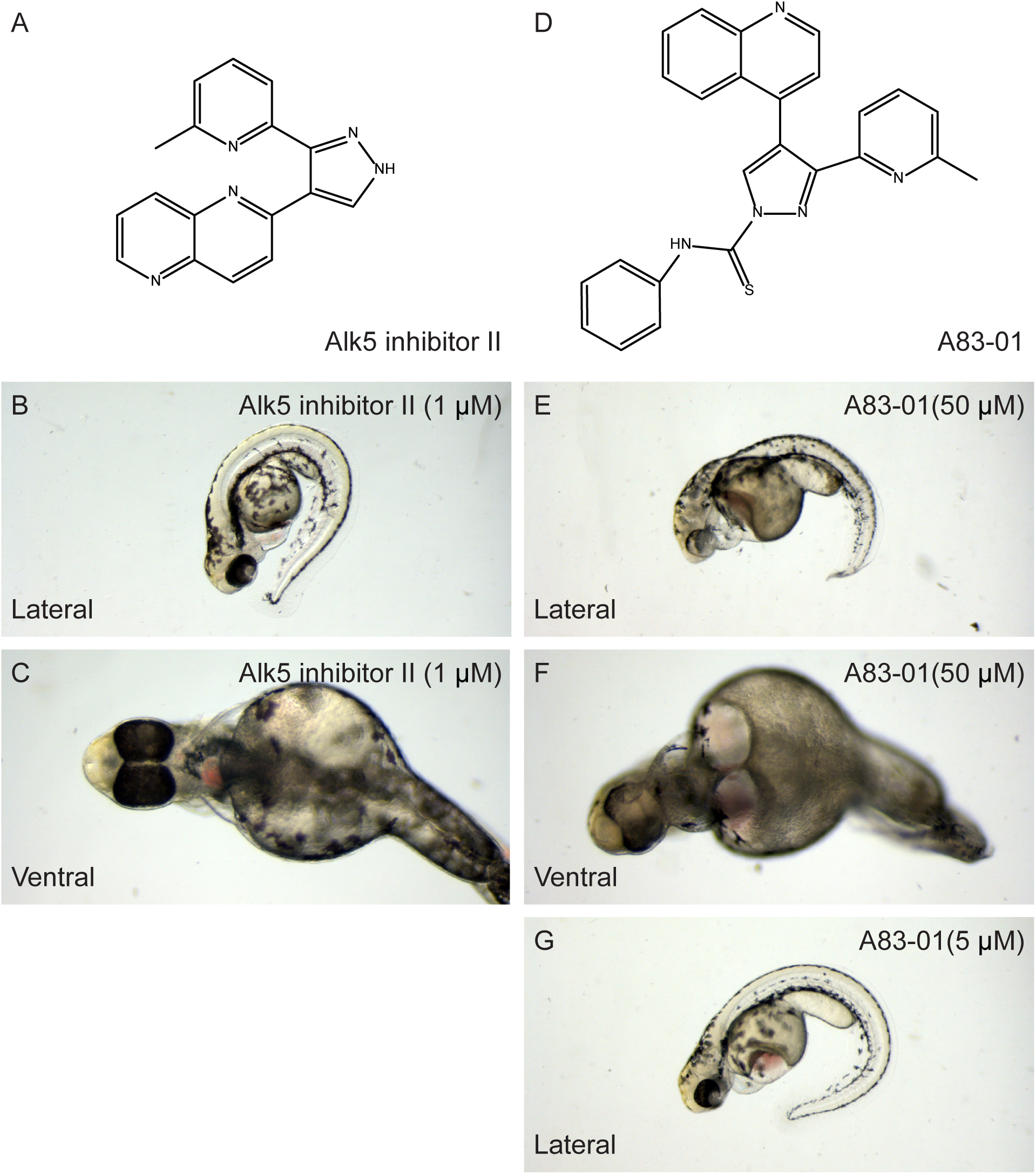
Inhibition of Alk5 using two independent Alk5 inhibitors induced developmental defects that were distinct from the developmental defects induced by known BMP inhibitors and cercosporamide.

## References

Aleström, P., D’Angelo, L., Midtlyng, P. J., Schorderet, D. F., Schulte-Merker, S., Sohm, F. and Warner, S. (2019). Zebrafish: Housing and husbandry recommendations. Lab. Anim. 0, 1–12.

Altman, J. K., Szilard, A., Konicek, B. W., Iversen, P. W., Glaser, H., Sassano, A., Vakana, E. and Graff, J. R. (2018). Inhibition of Mnk kinase activity by cercosporamide and suppressive effects on acute myeloid leukemia precursors.

Bauer, H., Lele, Z., Rauch, G. J., Geisler, R. and Hammerschmidt, M. (2001). The type I serine/threonine kinase receptor Alk8/Lost-a-fin is required for Bmp2b/7 signal transduction during dorsoventral patterning of the zebrafish embryo. Development 128, 849–858.

Bier, E. and De Robertis, E. M. (2015). BMP gradients: A paradigm for morphogen-mediated developmental patterning. Science 348,. aaa5838.

Cannon, J. E., Upton, P. D., Smith, J. C. and Morrell, N. W. (2010). Intersegmental vessel formation in zebrafish: Requirement for VEGF but not BMP signalling revealed by selective and non-selective BMP antagonists. Br. J. Pharmacol. 161, 140–149.

Cheng, V., Dasgupta, S., Reddam, A. and Volz, D. C. (2019). Ciglitazone-a human PPARγ agonist-disrupts dorsoventral patterning in zebrafish. PeerJ 1–20.

Dal-Pra, S., Fürthauer, M., Van-Celst, J., Thisse, B. and Thisse, C. (2006). Noggin1 and Follistatin-like2 function redundantly to Chordin to antagonize BMP activity. Dev. Biol. 298, 514–526.

Dasgupta, S., Vliet, S. M., Kupsco, A., Leet, J. K., Altomare, D. and Volz, D. C. (2017). Tris(1,3-dichloro-2-propyl) phosphate disrupts dorsoventral patterning in zebrafish embryos. PeerJ 1–16.

den Hertog, J. (2005). Chemical Genetics : Drug Screens in Zebrafish. Biosci. Rep. 25, 289–297.

Dennler, S., Itoh, S., Vivien, D., Dijke, P. Ten, Huet, S. and Gauthier, J. M. (1998). Direct binding of Smad3 and Smad4 to critical TGFβ-inducible elements in the promoter of human plasminogen activator inhibitor-type 1 gene. EMBO J. 17, 3091–3100.

Derwall, M., Malhotra, R., Lai, C. S., Beppu, Y., Aikawa, E., Seehra, J. S., Zapol, W. M., Bloch, K. D. and Yu, P. B. (2012). Inhibition of bone morphogenetic protein signaling reduces vascular calcification and atherosclerosis. Arterioscler. Thromb. Vasc. Biol. 32, 613–622.

Fujii, M., Takeda, K., Imamura, T., Aoki, H., Sampath, T. K., Enomoto, S., Kawabata, M., Kato, M., Ichijo, H. and Miyazono, K. (1999). Roles of bone morphogenetic protein type I receptors and Smad proteins in osteoblast and chondroblast differentiation. Mol. Biol. Cell 10, 3801–3813.

Gebruers, E., Cordero-Maldonado, M. L., Gray, A. I., Clements, C., Harvey, A. L., Edrada-Ebel, R., De Witte, P. A. M., Crawford, A. D. and Esguerra, C. V (2013). A phenotypic screen in zebrafish identifies a novel small-molecule inducer of ectopic tail formation suggestive of alterations in non-canonical Wnt/PCP signaling. PLoS One 8, 1–14.

Gomez-Puerto, M. C., Iyengar, P. V., García de Vinuesa, A., ten Dijke, P. and Sanchez-Duffhues, G. (2019). Bone morphogenetic protein receptor signal transduction in human disease. J. Pathol. 247, 9–20.

Hao, J., Ho, J. N., Lewis, J. A., Karim, K. A., Daniels, R. N., Gentry, P. R., Hopkins, C. R., Lindsley, C. W. and Hong, C. C. (2010). In vivo structure - Activity relationship study of dorsomorphin analogues identifies selective VEGF and BMP inhibitors. ACS Chem. Biol. 5, 245–253.

Hill, C. S. (2016). Transcriptional control by the SMADs. Cold Spring Harb. Perspect. Biol. 8,.

Hocking, J. C., Famulski, J. K., Yoon, K. H., Widen, S. A., Bernstein, C. S., Koch, S., Weiss, O., Agarwala, S., Inbal, A., Lehmann, O. J., et al. (2018). Morphogenetic defects underlie Superior Coloboma, a newly identified closure disorder of the dorsal eye. PLoS Genet. 14, 1–28.

Hoeksma, J., Misset, T., Wever, C., Kemmink, J., Kruijtzer, J., Versluis, K., Liskamp, M. J., Boons, G. J., Heck, A. J. R., Boekhout, T., et al. (2019). A new perspective on fungal metabolites: identification of bioactive compounds from fungi using zebrafish embryogenenis as read-out. Sci. Rep. 1–16.

Hosoya, T., Ohsumi, J., Hamano, K., Ono, Y. and Miura, M. (2011). Method for producing cercosporamide - Patent No.: US 7,939,081 B2.

Kajimoto, H., Kai, H., Aoki, H., Uchiwa, H., Aoki, Y., Yasuoka, S., Anegawa, T., Mishina, Y., Suzuki, A., Fukumoto, Y., et al. (2015). BMP type i receptor inhibition attenuates endothelial dysfunction in mice with chronic kidney disease. Kidney Int. 87, 128–136.

Kaplan, F. S., Xu, M., Seemann, P., Connor, J. M., Glaser, D. L., Carroll, L., Delai, P., Fastnacht-Urban, E., Forman, S. J., Gillessen-Kaesbach, G., et al. (2009). Classic and atypical fibrodysplasia ossificans progressiva (FOP) phenotypes are caused by mutations in the bone morphogenetic protein (BMP) type I receptor ACVR1. Hum. Mutat. 30, 379–390.

Katagiri, T. and Watabe, T. (2016). Bone morphogenetic proteins. Cold Spring Harb. Perspect. Biol. 1–27.

Kimmel, C. B., Ballard, W. W., Kimmel, S. R., Ullmann, B. and Schilling, T. F. (1995). Stages of embryonic development of the zebrafish. Dev Dyn 203, 253–310.

Kishimoto, Y., Lee, K. H., Zon, L., Hammerschmidt, M. and Schulte-Merker, S. (1997). The molecular nature of zebrafish swirl: BMP2 function is essential during early dorsoventral patterning. Development 124, 4457–4466.

Konicek, B. W., Stephens, J. R., McNulty, A. M., Robichaud, N., Peery, R. B., Dumstorf, C. A., Dowless, M. S., Iversen, P. W., Parsons, S., Ellis, K. E., et al. (2011). Therapeutic inhibition of MAP kinase interacting kinase blocks eukaryotic initiation factor 4E phosphorylation and suppresses outgrowth of experimental lung metastases. Cancer Res. 71, 1849–1857.

Korchynskyi, O. and Ten Dijke, P. (2002). Identification and functional characterization of distinct critically important bone morphogenetic protein-specific response elements in the Id1 promoter. J. Biol. Chem. 277, 4883–4891.

Kramer, C., Mayr, T., Nowak, M., Schumacher, J., Runke, G., Bauer, H., Wagner, D. S., Schmid, B., Imai, Y., Talbot, W. S., et al. (2002). Maternally Supplied Smad5 Is Required for Ventral Specification in Zebrafish Embryos Prior to Zygotic Bmp Signaling. Dev. Biol. 250, 263–279.

LaBonty, M. and Yelick, P. C. (2018). Animal models of Fibrosyplasia Ossificans Progressiva. Dev Dyn 247, 279–288.

Lefort, S. and Maguer-Satta, V. (2020). Targeting BMP signaling in the bone marrow microenvironment of myeloid leukemia. Biochem. Soc. Trans. 48, 411–418.

Little, S. C. and Mullins, M. C. (2004). Twisted gastrulation promotes BMP signaling in zebrafish dorsal-ventral axial patterning. Development 131, 5825–5835.

Liu, Y., Sun, L., Su, X. and Guo, S. (2016). Inhibition of eukaryotic initiation factor 4E phosphorylation by cercosporamide selectively suppresses angiogenesis, growth and survival of human hepatocellular carcinoma. Biomed. Pharmacother. 84, 237–243.

Mintzer, K. A., Lee, M. A., Runke, G., Trout, J., Whitman, M. and Mullins, M. C. (2001). Lost-a-fin encodes a type I BMP receptor, Alk8, acting maternally and zygotically in dorsoventral pattern formation. Development 128, 859–869.

Mucha, B. E., Hashiguchi, M., Zinski, J., Shore, E. M. and Mullins, M. C. (2018). Variant BMP receptor mutations causing fibrodysplasia ossificans progressiva (FOP) in humans show BMP ligand-independent receptor activation in zebrafish. Bone 109, 225–231.

Oxtoby, E. and Jowett, T. (1993). Cloning of the zebrafish krox-20 gene (krx-20) and its expression during hindbrain development. Nucleic Acids Res. 21, 1087–1095.

Persson U, Izumi H, Souchelnytskyi S, Itoh S, Grimsby S, Engström U, Heldin CH, Funa K, ten Dijke, P. The L45 Loop in Type I Receptors for TGF-beta Family Members Is a Critical Determinant in Specifying Smad Isoform Activation. FEBS Lett. 434, 83–87

Pignolo, R. J. and Kaplan, F. S. (2018). Clinical staging of Fibrodysplasia Ossificans Progressiva (FOP). Bone 109, 111–114.

Saeed, O., Otsuka, F., Polavarapu, R., Karmali, V., Weiss, D., Davis, T., Rostad, B., Pachura, K., Adams, L., Elliott, J., et al. (2012). Pharmacological suppression of hepcidin increases macrophage cholesterol efflux and reduces foam cell formation and atherosclerosis. Arterioscler. Thromb. Vasc. Biol. 32, 299–307.

Sanvitale, C. E., Kerr, G., Chaikuad, A., Ramel, M. C., Mohedas, A. H., Reichert, S., Wang, Y., Triffitt, J. T., Cuny, G. D., Yu, P. B., et al. (2013). A New Class of Small Molecule Inhibitor of BMP Signaling. PLoS One 8, e62721.

Schmid, B., Fürthauer, M., Connors, S. A., Trout, J., Thisse, B., Thisse, C. and Mullins, M. C. (2000). Equivalent genetic roles for bmp7/snailhouse and bmp2b/swirl in dorsoventral pattern formation. Development 127, 957–967.

Shen, Q., Little, S. C., Meiqi, X., Haupt, J., Ast, C., Katagiri, T., Mundlos, S., Seemann, P., Kaplan, F. S., Mullins, M. C., et al. (2009). The fibrodysplasia ossificans progressiva R206H ACVR1 mutation activates BMP-independent chondrogenesis and zebrafish embryo ventralization. J. Clin. Invest. 119, 3462–3472.

Smith, K. A., Joziasse, I. C., Chocron, S., Van Dinther, M., Guryev, V., Verhoeven, M. C., Rehmann, H., Van Der Smagt, J. J., Doevendans, P. A., Cuppen, E., et al. (2009). Dominant-negative alk2 allele associates with congenital heart defects. Circulation 119, 3062–3069.

Sugawara, F., Takahashi, N., Strobel, S., Strobel, G., Larsen, R. D., Berglund, D. L., Gray, G., Coval, S. J., Stout, T. J. and Clardy, J. (1991). The Structure and Biological Activity of Cercosporamide from Cercosporidium henningsii. J. Org. Chem. 56, 909–910.

Sussman, A., Huss, K., Chio, L., Heidler, S., Shaw, M., Ma, D., Zhu, G., Campbell, R. M., Park, T., Kulanthaivel, P., et al. (2004). Discovery of Cercosporamide, a Known Antifungal Natural Product, as a Selective Pkc1 Kinase Inhibitor through High-Throughput Screening. 3, 932–943.

Suzuki, T., Nakano, M., Komatsu, M., Takahashi, J., Kato, H. and Nakamura, Y. (2020). ZMAT2, a newly-identified potential disease-causing gene in congenital radioulnar synostosis, modulates BMP signaling. Bone 136, 115349.

Taylor, K. R., Vinci, M., Bullock, A. N. and Jones, C. (2014). ACVR1 mutations in DIPG: Lessons learned from FOP. Cancer Res. 74, 4565–4570.

Thisse, C. and Thisse, B. (2008). High-resolution in situ hybridization to whole-mount zebrafish embryos. Nat. Protoc. 3, 59–69.

Weinberg, E. S., Allende, M. L., Kelly, C. S., Abdelhamid, A., Murakami, T., Andermann, P., Doerre, O. G., Grunwald, D. J. and Riggleman, B. (1996). Developmental regulation of zebrafish MyoD in wild-type, no tail and spadetail embryos. Development 122, 271–280.

Wiley, D. S., Redfield, S. E. and Zon, L. I. (2017). Chemical screening in zebrafish for novel biological and therapeutic discovery. Methods Cell Biol. 138, 651–679.

Yang, Y. and Thorpe, C. (2011). BMP and non-canonical Wnt signaling are required for inhibition of secondary tail formation in zebrafish. Development 138, 2601–2611.

Ye, M., Berry-Wynne, K. M., Asai-Coakwell, M., Sundaresan, P., Footz, T., French, C. R., Abitbol, M., Fleisch, V. C., Corbett, N., Allison, W. T., et al. (2009). Mutation of the bone morphogenetic protein GDF3 causes ocular and skeletal anomalies. Hum. Mol. Genet. 19, 287–298.

Yu, P. B., Hong, C. C., Sachidanandan, C., Babitt, J. L., Deng, D. Y., Hoyng, S. A., Lin, H. Y., Bloch, K. D. and Peterson, R. T. (2008). Dorsomorphin inhibits BMP signals required for embryogenesis and iron metabolism. Nat. Chem. Biol. 4, 33–41.

Zhou, G., Myers, R., Li, Y., Chen, Y., Shen, X., Fenyk-melody, J., Wu, M., Ventre, J., Doebber, T., Fujii, N., et al. (2001). Role of AMP-activated protein kinase in mechanism of metformin action. J. Clin. Invest. 108, 1167–1174.

